# Defining the transmissible dose 50%, the donor inoculation dose that results in airborne transmission to 50% of contacts, for two pandemic influenza viruses in ferrets

**DOI:** 10.1101/2025.03.04.641289

**Authors:** C. J. Field, K.M. Septer, D.R. Patel, V.C Weaver, D.G. Sim, R. H. Restori, T.C. Sutton

## Abstract

Ferrets are widely used to model airborne transmission of influenza viruses in humans. Airborne transmission is evaluated by infecting donor ferrets with a high virus dose (10^6^ infectious units) and monitoring transmission to contact animals sharing the same airspace. However, humans can be infected with a broad range of influenza virus doses. Therefore, we evaluated the relationship between virus inoculation dose and transmission for two pandemic influenza viruses in ferrets. Donor ferrets were inoculated with 10^0^ to 10^6^ tissue culture infectious dose 50 (TCID_50_) of the 2009 pandemic H1N1 or 1968 H3N2 pandemic virus, and were then paired with respiratory contacts. Using the proportion of donors that became infected across virus doses, we calculated the infectious dose 50 (ID_50_). Subsequently, by comparing the proportion of respiratory contacts that became infected, we calculated the transmissible dose 50% (TD_50_): the donor inoculation dose that resulted in transmission to 50% of contacts. For the 2009 H1N1 virus, the ID_50_ and TD_50_ were equivalent at <1 TCID_50_. However, for the 1968 H3N2 virus, the ID_50_ and TD_50_ were <4 and 10^3.49^ TCID_50_, respectively. The increased TD_50_ for the H3N2 virus was associated with reductions in peak viral titers and viral shedding in donors over decreasing virus inoculation doses. Collectively, these studies define a new measure of transmission that permits comparisons of transmissibility over a log scale. Using this metric, we show the 1968 pandemic H3N2 virus has reduced transmissibility in ferrets relative to the 2009 pandemic H1N1 virus.

**Importance:** Ferrets are the gold standard animal model used to assess the transmissibility of influenza viruses. Airborne transmission is evaluated by infecting donor ferrets with a high virus dose and monitoring transmission to contact animals sharing the same airspace. However, the relationship between inoculation dose and transmission has not been evaluated in ferrets. Therefore, we performed studies evaluating airborne transmission of the 2009 pandemic H1N1 and 1968 pandemic H3N2 viruses over log scale reductions in donor inoculation doses. Using the results of these studies, we define a new measure of transmission, the transmissible dose 50%, which is the donor inoculation dose at which a virus is transmitted to 50% of contacts. Importantly, this metric permits the evaluation of transmissibility over a 10-100 fold scale, and using this metric, we demonstrate the 1968 pandemic H3N2 virus has reduced transmissibility compared to the 2009 pandemic H1N1 virus in ferrets.

## Introduction

Influenza A viruses cause annual epidemics and sporadic pandemics. Currently, influenza A viruses of the H1N1 and H3N2 subtypes co-circulate in humans. Each year during seasonal influenza epidemics, these viruses infect up to 1 billion people worldwide, leading to 3-5 million cases of severe disease, and 250-650,000 deaths (1–3). Influenza pandemics occur at irregular intervals when a virus that is antigenically novel to humans spills over from an animal reservoir (*i.e.* waterfowl or pigs) and transmits efficiently from person-to-person via the airborne route (4). During influenza pandemics, the disease burden exceeds that of annual epidemics, and disease severity can be severe (5, 6). To study influenza virus pathogenesis and transmission, several animal models have been developed (7–9). Of these models, the ferret is the only animal model that recapitulates both disease and airborne transmission observed in humans. As a result, ferrets are the predominant animal model used to study transmission, and when gauging the pandemic risk posed by an emerging virus, both the WHO and CDC utilize data on the transmissibility of the virus in ferrets as part of this assessment (10, 11).

To study airborne transmission in ferrets, donor animals are typically inoculated with 10^6^ infectious units of a virus. Twenty-four hours post-infection, each donor (DR) animal is introduced into a transmission cage with a respiratory contact (RC) ferret. The transmission cages are designed such that the DR and RC share the same airspace but cannot have direct physical contact. Nasal wash samples are then collected from both the DR and RC over 10-14 days, and the samples are assayed for infectious virus. On day 14 or 21 post-pairing, blood samples are also collected from both the DR and RC to assess seroconversion (7, 8, 12). RCs are considered infected if they shed infectious virus and/or seroconvert. Transmission studies are often performed using 3 or 4 DR:RC pairs, and the transmission efficiency of a virus is expressed as a percentage of the contacts that become infected (*i.e.* 2 of 4 RCs infected equals 50% transmission efficiency). Based on studies with highly transmissible human seasonal and pandemic viruses, and poorly transmissible avian influenza viruses, viruses that transmit to 66% or greater of RCs are considered to transmit efficiently via the airborne route in ferrets and likely to have some degree of transmissibility in humans. While viruses that transmit to less than 66% of RCs are considered to have inefficient transmission in ferrets and unlikely to exhibit transmissibility in humans (13).

Human challenge studies have found that humans can be infected with influenza viruses over a wide range of inoculation doses (reviewed in(14)). While most challenge studies use high doses between 10^4^-10^7^ infectious units to ensure individuals become infected, studies evaluating the human infectious dose 50%, especially via aerosol inoculation, have found as little as 0.6-5.0 tissue culture infectious dose 50s (TCID_50_) can establish an infection in the nose of volunteers (15). However, despite the extensive use of ferrets to study influenza virus transmission, the impact of inoculation dose on transmission has not been evaluated. Therefore, we conducted studies to define the relationship between infectious dose and airborne transmission in ferrets for two pandemic influenza viruses: the 1968 pandemic H3N2 virus (A/Hong Kong/1/1968) [1968 H3N2] and the 2009 pandemic H1N1 (A/California/07/2009 (H1N1pdm09)) [2009 H1N1] virus. To minimize the role of passage history, viruses were generated by reverse genetics and minimally passaged in Madin-Darby Canine Kidney (MDCK) cells. DR ferrets were inoculated with 10-100-fold decreasing doses of virus, and transmission to RCs was monitored over 14 days. By determining the infection status of the DRs over log-scale inoculation doses, we calculated the ferret infectious dose 50% (ID_50_). Subsequently, using the proportion of RCs that became infected at each virus dose, we calculated the DR inoculation dose that resulted in transmission to 50% of contacts. We designated this dose the transmissible dose 50% (TD_50_). In DR ferrets, the 2009 H1N1 and 1968 H3N2 viruses both had low ID_50_ values. As the donor inoculation dose was reduced, the 2009 H1N1 virus retained its transmissibility and had a TD_50_ equivalent to its ID_50_. In contrast, the 1968 H3N2 virus exhibited reduced transmission over decreasing inoculation doses leading to an increase in the TD_50_. This indicated the 1968 H3N2 virus has reduced transmissibility in ferrets relative to the 2009 H1N1 virus.

## Results

### The 2009 pandemic H1N1 virus transmits efficiently to respiratory contact ferrets over decreasing donor inoculation doses

To evaluate the relationship between infectious dose and transmission, DR ferrets (n=4/inoculation dose) were inoculated with 10^6^, 10^4^, 10^2^, 10^1^, or 10^0^ TCID_50_ of recombinant 2009 H1N1 virus. Twenty-four hours post-infection, each DR was introduced into an airborne transmission cage with a paired RC. Within the transmission cage, the animals were oriented such that airflow was directional from the DI to the RC. Nasal wash samples were then collected from the ferrets every other day for 14 days, and serum was collected from DRs and RCs on day 21 post-inoculation of the DR.

During the 14-day sampling period, weight loss and clinical signs were monitored in both the DR and RC animals. Clinical signs were mild in all the DRs and there were no significant differences in weight loss and changes in body temperatures across virus inoculation doses. When RC animals became infected, clinical disease was also limited and no significant differences were observed across DR inoculation doses (**Table 1**). For these studies, we used stringent criteria to define an RC as infected, and this included virus shedding in the nasal wash on at least 1 day and seroconversion. In the DR animals, infectious virus was detected in the nasal wash by 1 day post-inoculation (dpi) for doses of 10^6^ and 10^4^ TCID_50_, and by 3 dpi for 10^2^, 10^1^, and 10^0^ TCID_50_ (**Fig 1**). While not significantly different across doses, the start of viral shedding was delayed by an average of 1.5 days from the highest to lowest inoculation dose (**Fig 1F**). All (100%) of DRs shed infectious virus in the nasal wash for all virus doses except the lowest dose of 10^0^ TCID_50_ where one DR did not shed virus. In the infected DRs, all animals cleared the virus by 11 dpi.

**Figure 1.**
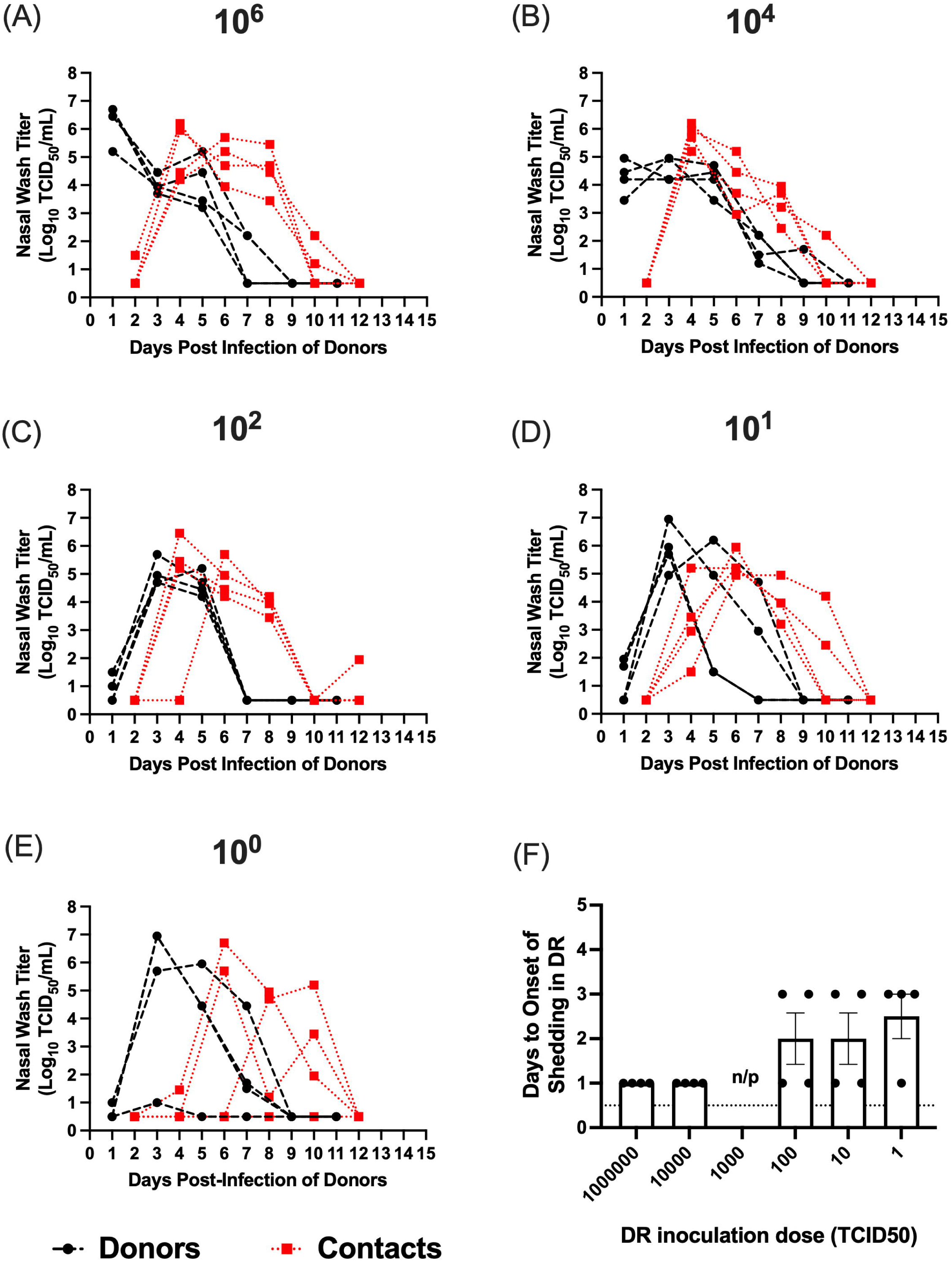
Airborne transmission of the 2009 H1N1 virus over decreasing donor inoculation doses. Shown in (A-E) are nasal wash titers in donors (black lines) and respiratory contacts (red lines) for decreasing donor inoculation doses. Donor inoculation dose in TCID_50_ is shown above each a panel. Donor ferrets were inoculated with different doses of virus in a 1 mL volume, and 24 hours post-infection, these animals were paired with respiratory contacts. Nasal wash samples were then collected from donors and contacts on alternating days. (F) shows the average number of days +/-standard error to onset of viral shedding in the DRs across virus inoculation doses. Note: in panels (D) and (E) two donors have viral shedding curves that are superimposed.

**Table 1.** Clinical signs for DR and RC ferrets infected with the 2009 H1N1 and 1968 H3N2 viruses.

| Inoculation Dose | Donors |  | Respiratory Contacts |  |
| --- | --- | --- | --- | --- |
|  | Maximum Change in Body Temperature (°C)<sup>a</sup><br>(mean +/- SE) | Maximum Weight Loss (%)<sup>a</sup><br>(mean +/- SE) | Maximum Change in Body Temperature (°C)<sup>a</sup><br>(mean +/- SE) | Maximum Weight Loss (%)<sup>a</sup><br>(mean +/- SE) |
| **A/California/07/2009 (H1N1pdm09)** |  |  |  |  |
| **10<sup>6</sup>** | 0.80 +/-0.2 | 7.89 +/- 1.5 | 0.93 +/- 0.3 | 3.37+/- 1.3 |
| **10<sup>4</sup>** | 0.48 +/-0.1 | 6.84 +/- 1.7 | 0.33 +/-0.1 | 6.10 +/- 2.3 |
| **10<sup>3</sup>** | n/p | n/p | n/p | n/p |
| **10<sup>2</sup>** | 0.75 +/-0.5 | 6.30 +/- 2.1 | 1.63 +/- 0.3 | 8.87+/-1.17 |
| **10<sup>1</sup>** | 0.35 +/-0.2 | 8.18 +/- 1.6 | 0.93 +/- 0.3 | 8.13+/-1.17 |
| **10<sup>0</sup>** | 0.45+/- 0.1 | 7.17 +/- 2.7 | 0.70+/-0.3 | 8.63+/- 1.00 |
| **A/Hong Kong/1/1968 (H3N2)** |  |  |  |  |
| **10<sup>6</sup>** | 0.54 +/-0.1 | 6.77 +/-2.2 | 0.62 +/- 0.4 | 6.05 +/- 2.8 |
| **10<sup>4</sup>** | 0.48 +/-0.3 | 9.53 +/- 4.5 | 0.60 +/- 0.2 | 7.71 +/- 1.2 |
| **10<sup>3</sup>** | 0.64 +/-0.2 | 1.74 +/- 1.0 | 0.10<br>(only 1 RC infected) | 0<br>(only 1 infected RC and animal gained weight) |
| **10<sup>2</sup>** | 0.85 +/-0.3 | 18.6 +/- 5.9<sup>b</sup> | 1.00 +/- 0.1<br>(2 RCs infected) | 10.8+/- 10.8<br>(2 RCs infected) |
| **10<sup>1</sup>** | 0.18 +/-0.1 | 1.23 +/- 1.0 | No data<sup>†</sup> | 1.78<br>(only 1 RC infected) |
| **10<sup>0</sup>** | 0.43 +/-0.2 | 5.27 +/- 2.6 | 0.60 +/- 0.3 | 2.53 +/- 0.5 |

Notes:
<sup>a</sup>uninfected DR and RC animals were excluded from calculations of average temperature change and weight loss except for A/Hong Kong/1/1968 (H3N2) at dose of 10<sup>0</sup> as no DR or RCs were infected. This later data is included for comparison of temperature changes and weight loss at other doses and between viruses.
<sup>b</sup>water bottles were malfunctioning resulting in weight loss
<sup>†</sup> No temperature data as only one RC was infected and its temperature microchip malfunction.

In the RCs, at donor inoculation doses of 10^6^, 10^4^, 10^2^, and 10^1^ TCID_50_, all the animals became infected, and at a DR inoculation dose of 10^0^ TCID_50_, 3 of 4 (75%) of the RCs became infected. This was confirmed by serology and all RC ferrets that shed virus in the nasal wash seroconverted by 21 dpi of the DRs (**Table 2**). Infectious virus was detected in nasal washes from RCs by 4 dpi of the DRs in the 10^6^ and 10^4^ TCID_50_ groups, and by 4-8 dpi in the 10^2^, 10^1^, and 10^0^ TCID_50_ groups. All RC ferrets stopped shedding virus in the nasal wash by 12 dpi of the DRs.

**Table 2.**
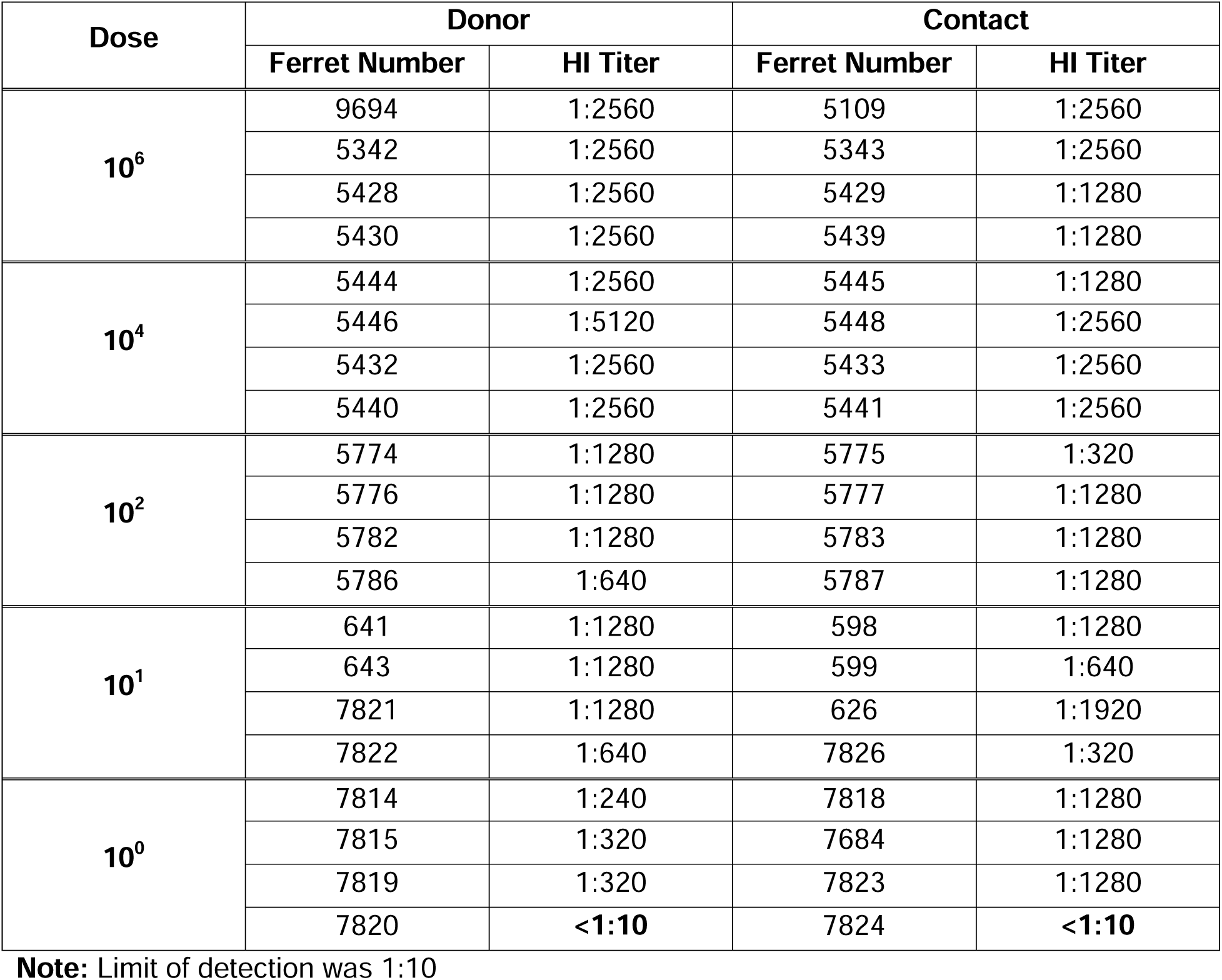
Hemagglutination inhibition antibody titers in DR and RC ferrets for airborne transmission studies with the 2009 H1N1 virus.

Using the number of infected DRs at each inoculation dose, we calculated the ID_50_ for the 2009 H1N1 virus. As all the DRs became infected at virus inoculations doses of 10^1^ TCID_50_ and higher, and 75% of the DRs were infected at an inoculation dose of 10^0^ TCID_50_, the ID_50_ for the 2009 H1N1 virus was <1 TCID_50_. Using the number of RCs infected at each DR inoculation dose, we then calculated the TD_50_ which was also <1 TCID_50_ (**Table 4**). Therefore, the 2009 H1N1 virus was highly transmissible even at very low inoculation doses, and the ID_50_ and TD_50_ were comparable.

### The 1968 H3N2 virus exhibited variable and reduced airborne transmission to contacts over decreasing donor inoculation doses

Next, we evaluated the relationship between inoculation dose and transmission for the 1968 H3N2 virus. DR ferrets were intranasally inoculated with 10^6^ to 10^0^ TCID_50_ of the A/Hong Kong/1/1968 (H3N2) [H3N2 1968] virus. After performing initial studies, transmission studies at doses of 10^3^ and 10^6^ were repeated to verify our findings. Consistent with our findings for the 2009 H1N1 virus, during the 14-day sampling period, we did not observe any significant differences in the clinical signs among infected DRs at different inoculation doses or in the infected RC animals (**Table 1**, note: at inoculation dose of 10^2^, DRs lost weigh due to malfunctioning water bottles). In the DRs, infectious virus was detected in the nasal wash by 1-3 dpi at the 10^6^ and 10^4^ TCID_50_ inoculation doses, and by 3-5 dpi for 10^3^, 10^2^, and 10^1^ TCID_50_ doses (**Fig 2**). On average, DRs took 2 days longer to start shedding virus in the nasal wash at an inoculation dose of 10^1^ compared to 10^6^ TCID_50_, and there was a significant delay in viral shedding for DRs in the 10^2^ and 10^1^ groups relative to the 10^6^ group (**Fig 2I**). All (100%) of the DRs became infected at inoculation doses of 10^6^-10^1^ TCID50 and these animals also seroconverted; however, no donors inoculated with 10^0^ TCID50 shed virus at any time point and these animals did not seroconvert (**Fig 2** and **Table 3**). All infected DRs cleared the virus by 11 dpi regardless of inoculation dose.

**Figure 2.**
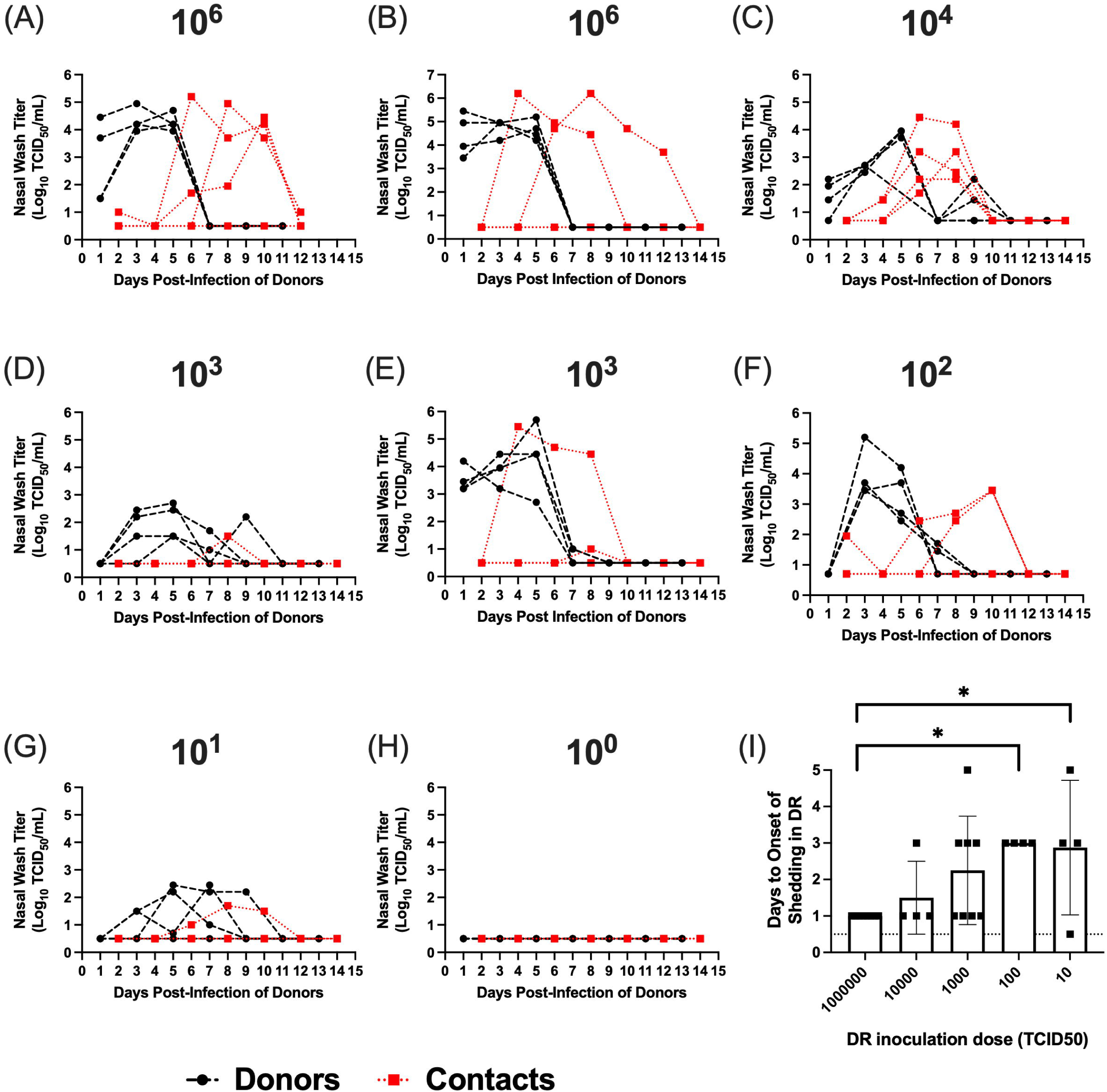
Airborne transmission of the 1968 H3N2 virus over decreasing donor inoculation doses. Shown in (A-H) are nasal wash titers in donors (black lines) and respiratory contacts (red lines) for transmission studies using different donor inoculation doses. Donor inoculation dose in TCID_50_ is shown above each a panel, and for the 10^6^ and 10^3^ doses two replicate studies were performed (panel B and E). Donor ferrets were inoculated with different doses of virus in a 1 mL volume, and 24 hours post-infection, these animals were paired with respiratory contacts. Nasal wash samples were then collected from donors and contacts on alternating days. (I) shows the average number of days +/-standard error to onset of viral shedding in the DRs across virus inoculation doses. Note: in panel (F), two donors have viral shedding curves that are superimposed.

**Table 3.** Hemagglutination inhibition antibody titers in DR and RC ferrets for airborne transmission studies with the 1968 H3N2 virus.

| Dose | Donor |  | Contact |  |
| --- | --- | --- | --- | --- |
|  | Ferret Number | HI Titer | Ferret Number | HI Titer |
| **10<sup>6</sup>** | 1401 | 1:2560 | 1408 | 1:2560 |
|  | 2120 | 1:640 | 2124 | 1:1280 |
|  | 2118 | 1:2560 | 2119 | **<1:10** |
|  | 2103 | 1:2560 | 2104 | 1:2560 |
|  | 4627 | 1:1280 | 4634 | **<1:10** |
|  | 4629 | 1:1280 | 4632 | 1:1280 |
|  | 4617 | 1:640 | 4620 | **<1:10** |
|  | 4619 | 1:320 | 4622 | 1:640 |
| **10<sup>4</sup>** | 6321 | 1:160 | 6322 | 1:640 |
|  | 6323 | 1:320 | 6324 | 1:640 |
|  | 6333 | 1:160 | 6334 | 1:320 |
|  | 6335 | 1:320 | 6336 | 1:640 |
| **10<sup>3</sup>** | 7436 | 1:640 | 7439 | **<1:10** |
|  | 7440 | n/p | 7441 | **<1:10** |
|  | 7420 | 1:1280 | 7425 | **<1:10** |
|  | 7428 | 1:160 | 7429 | **<1:10** |
|  | 4623 | 1:640 | 4633 | **<1:10** |
|  | 4625 | 1:1280 | 4624 | **<1:10** |
|  | 4613 | 1:1280 | 4614 | **<1:10** |
|  | 4615 | 1:1280 | 4618 | 1:1280 |
| **10<sup>2</sup>** | 6313 | 1:320 | 6314 | **<1:10** |
|  | 671 | 1:1280 | 626 | 1:640 |
|  | 6327 | 1:320 | 6328 | **<1:10** |
|  | 6329 | 1:1280 | 6330 | 1:640 |
| **10<sup>1</sup>** | 11 | 1:640 | 7430 | **<1:10** |
|  | 7433 | 1:320 | 7434 | 1:640 |
|  | 7423 | 1:320 | 7426 | **<1:10** |
|  | 7427 | 1:480 | 007 | **<1:10** |
| **10<sup>0</sup>** | 5 | **<1:10** | 7442 | **<1:10** |
|  | 7437 | **<1:10** | 7443 | **<1:10** |
|  | 7418 | **<1:10** | 7422 | **<1:10** |
|  | 7419 | **<1:10** | 7424 | **<1:10** |

**Note:** Limit of detection was 1:10. n/p denotes not performed. Serum sample was lost during processing.

**Table 4.**
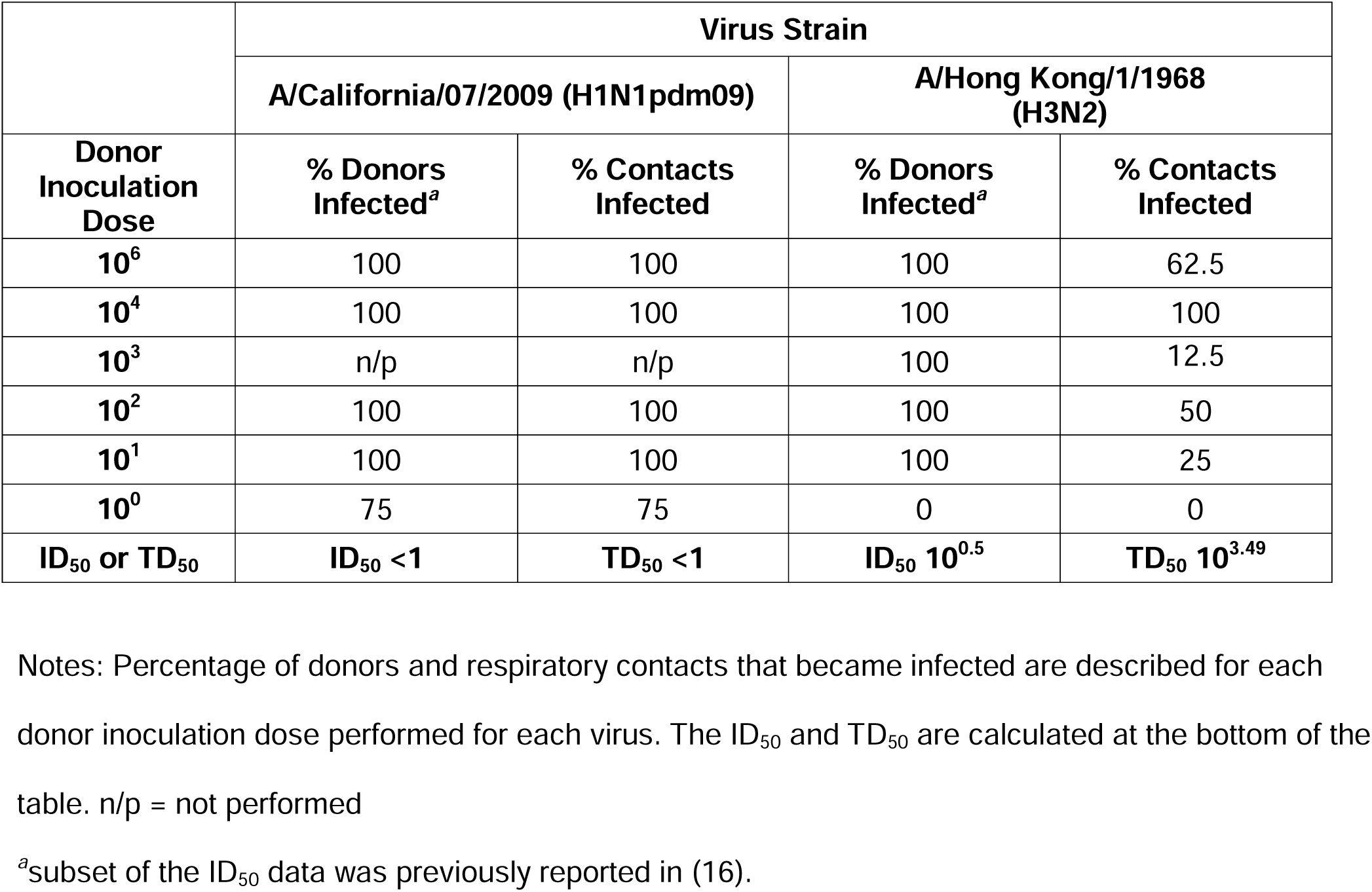
Summary of airborne transmission studies in ferrets using the 2009 H1N1 and 1968 H3N2 viruses.

At an inoculation dose of 10^6^ TCID_50_, 3 of 4 (75%), and 2 of 4 (50%) RCs shed infectious virus and seroconverted in the two replicate studies, respectively (**Fig 2A and B**). Thus, a total of 5 of 8 RCs (62.5%) became infected at a DR inoculation dose of 10^6^. When DRs were inoculated with 10^4^ TCID_50_ of virus, all 4 RCs (100%) shed virus and seroconverted (**Table 3**). This was the only inoculation dose for the 1968 H3N2 virus where the virus was transmitted to all the RCs. For the replicate studies with a DR inoculation dose of 10^3^, 2 of 4, and 1 of 4 RCs shed infectious virus in each replicate, respectively; however, neither RC in the first replicate seroconverted and these animals only shed low titers of virus on a single day. In the second replicate, the single RC that shed virus also seroconverted. Thus, based on our criteria, a total of 1 of 8 RCs became infected (12.5%) (**Figure 2D-E** and **Table 3**). When the DR inoculation dose was further reduced to 10^2^ and 10^1^ TCID_50_, 50% and 25% of RCs shed virus and seroconverted, respectively (**Table 3**). Virus was cleared from the nasal wash by 12 dpi for all RCs. At a DR inoculation dose of 10^0^ TCID_50_, none of the RCs became infected consistent with a lack of viral shedding from their DRs.

Using the criteria of viral shedding combined with seroconversion to define an animal as infected, we calculated the ID_50_ and TD_50_ for the 1968 H3N2 virus. The ID_50_ in DRs was 10^0.5^ TCID_50_, and the TD_50_ in RCs was 10^3.49^ TCID_50_. This ID_50_ is similar to the 2009 H1N1 virus; however, the TD_50_ is 3 orders of magnitude higher (**Table 4**). Therefore, while both the 2009 H1N1 and 1968 H3N2 viruses have low ID_50_’s, these viruses exhibit differing degrees of transmissibility, with the 1968 H3N2 virus having more variable and reduced transmission at low inoculation doses in ferrets.

#### The 2009 H1N1 virus replicated to similar peak titers across inoculation doses while the 1968 H3N2 virus exhibited reduced replication and overall shedding at lower inoculation doses

To determine if differences in the TD_50_ were associated with differences in viral replication kinetics, we compared peak titers and total viral shedding in the DRs across inoculation doses for the 2009 H1N1 and 1968 H3N2 viruses. We found DRs infected with the 2009 H1N1 virus had significantly higher peak titers than DRs infected with H3N2 virus at inoculation doses of 10^6^, 10^4^, 10^1^, and 10^0^ TCID_50_ (**Fig 3A**). Moreover, peak titers for the 2009 H1N1 virus were similar over decreasing virus inoculation doses. In contrast, peak titers in DRs infected with the 1968 H3N2 virus were reduced at lower inoculation doses. Peak titers for the 1968 H3N2 10^1^ TCID_50_ group where significantly lower than those for animals inoculated with 10^6^ TCID_50_, and DRs inoculated with 10^0^ TCID_50_ did not become infected. To quantify total virus shedding, we analyzed area under the curve of viral titers for each infected DR. For all inoculation doses, DRs infected with the 2009 H1N1 virus shed more virus than those infected with the H3N2 virus (**Fig 3B**). This was statistically significant for inoculation doses of 10^4^, 10^1^, and 10^0^ TCID_50_. There was also a significant reduction in the total amount of virus shed in DRs infected with 10^1^ TCID_50_ of the H3N2 virus relative to those infected with 10^6^ TCID_50_, and no ferrets shed virus at the 10^0^ TCID_50_ dose. Therefore, the higher TD_50_ for 1968 H3N2 virus was associated with reduced peak titers and virus shedding compared to the 2009 H1N1 virus, although, these metrics were not significantly different at all doses tested.

**Figure 3.**
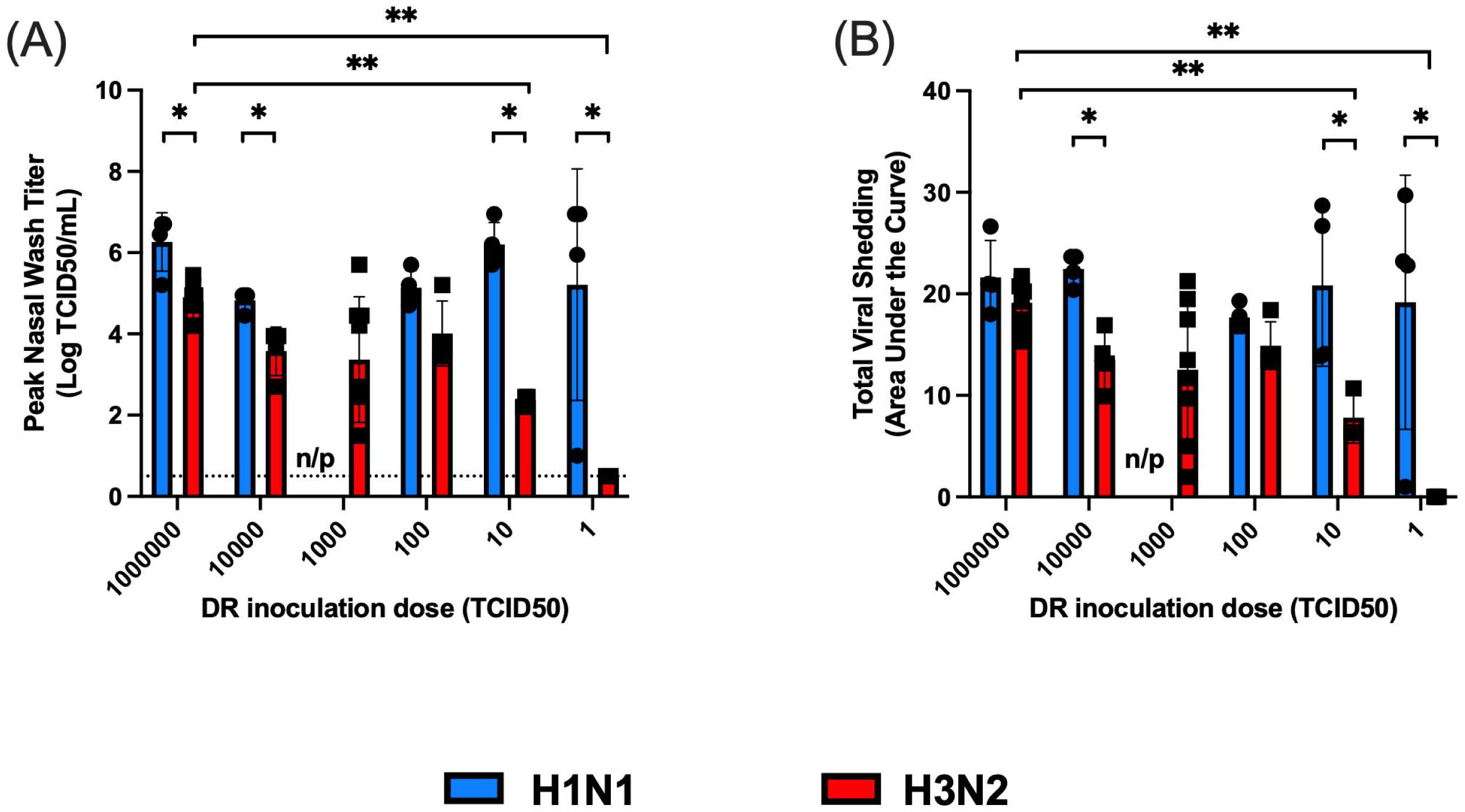
Analysis of peak viral titers and total viral shedding in donor ferrets across decreasing virus inoculation doses for the 2009 H1N1 and 1968 H3N2 viruses. Shown in (A) and (B) are peak viral titers and total virus shedding, respectively, for DR ferrets infected with the 2009 H1N1 or 1968 H3N2 viruses over decreasing virus inoculation doses. Blue and red bars represent the 2009 H1N1 and 1968 H3N2 viruses, respectively. Total viral shedding was determined via area under the curve for the nasal wash titer. Peak titers and total viral shedding were compared across virus doses for a given virus strain, and between virus strains for a specific virus dose. *significant differences between virus strains at the indicated dose (p<0.05). **significant differences between virus doses for virus strain (p<0.05). n/p denotes not performed because this virus dose was not tested in ferrets for the 2009 H1N1 virus.

## Discussion

The ferret model of human influenza virus transmission typically utilizes a high donor inoculation dose; however, humans are susceptible to influenza virus infection over a wide range of doses. To address this gap, we conducted studies to determine the relationship between inoculation dose and airborne transmission in ferrets. We inoculated DR ferrets with 10-100-fold decreasing doses of the 2009 H1N1 or 1968 H3N2 virus and assessed transmission to RCs. In the DR ferrets, both the 2009 H1N1 and 1968 H3N2 viruses had a low ID_50_ of <1 and 10^0.5^ TCID50, respectively. We previously published a subset of this ID_50_ data as part of our risk assessment studies on an H5N1 virus isolated from mink (16). Here, we build upon these studies and assessed transmission from directly inoculated ferrets to respiratory contact animals. We found that despite both viruses having a low ID_50_, the viruses differed in their transmissibility. To capture differences in transmissibility, we defined a new metric, the transmissible dose 50%: the DR inoculation dose that results in transmission to 50% of RCs. We found the 2009 H1N1 virus had a low TD_50_ of <1 TCID_50_ which is comparable to the ID_50_ for this virus. This indicates that if a DR ferret becomes infected with the 2009 H1N1 virus, it is likely to transmit the virus to its RC. In contrast, the TD_50_ for the 1968 H3N2 virus was several orders of magnitude higher and was 10^3.49^ TCID_50_. Therefore, for the 1968 H3N2 virus, much higher inoculation doses are required to consistently support airborne transmission.

To assess the relationship between viral shedding and transmission, we then analyzed peak titers and total viral shedding across DR inoculation doses. We found the 2009 H1N1 virus consistently replicated to comparable peak titers and was shed at similar levels regardless of inoculation dose. In contrast, there were reductions in peak titers and total virus shedding for the 1968 H3N2 virus as DR inoculation dose was reduced. Interestingly, for the 1968 H3N2 virus, we observed all the RC animals became infected with a DR inoculation dose of 10^4^ TCID_50,_ while reductions in transmission were observed at higher and lower DR inoculation doses. As high inoculation doses could potentially stimulate a robust innate immune response and lower doses may not support high levels of replication, this suggests for the H3N2 virus there may be an optimal inoculation dose that supports efficient transmission. This requires further evaluation using other representative 1968 H3N2 pandemic strains and more contemporary H3N2 viruses.

Our findings that ferrets can be infected with low doses of the 2009 H1N1 and 1968 H3N2 viruses are consistent with those reported by other groups (17–20). When the ID_50_ was determined in ferrets for several swine H1N1 reassortant viruses and the 2009 pandemic H1N1 virus, the ID_50_ for these viruses varied from <10 to 18 TCID_50_ (17). Our findings are also consistent with ferret studies evaluating replication of the A/California/04/2009 (H1N1pdm09) isolate at reduced inoculation doses: 10^6^, 10^4^, and 10^2^ TCID_50_ (18). At each virus dose, all ferrets became infected and peak viral replication in animals inoculated with 10^2^ TCID_50_ was delayed. Moreover, across the 3 inoculation doses, the A/California/04/2009 (H1N1pdm09) virus replicated to similar peak titers. In separate studies, ferrets were infected with 10^2^ or 10^5^ TCID_50_ of the A/California/07/2009 (H1N1pdm09) isolate, and replication kinetics were monitored over 9 days. Consistent with our findings at an inoculation dose 10^4^ TCID_50_ (**Fig 1B**), inoculation with 10^5^ TCID_50_ resulted in a plateau in virus titers from days 1-5, followed by a decline in titers on days 7 and 9. At an inoculation dose of 10^2^ TCID_50_, replication kinetics mirrored those of our studies (**Fig 1C**). Shortly after virus inoculation there was minimal viral shedding, followed by replication to peaks titers between 3-5 dpi, and then viral clearance by 9 dpi (19).

In studies with the seasonal A/Victoria/3/1975 (H3N2) virus, replication kinetics were analyzed in pairs of ferrets infected with 10^4^, 10^2^, and 10^1^ pfu of virus. Based on these studies, the ID_50_ was determined to be 10^0.66^ or 5 plaque forming units (pfu). This is directly comparable to the ID_50_ for the 1968 H3N2 virus. At the lower inoculation doses of 10^2^ and 10^1^, viral titers of the seasonal H3N2 virus peaked on day 2 or 3, and then declined steadily over the next 5 days. At an inoculation dose of 10^4^ pfu, viral titers peaked on day 2, declined to near the limit of detection on day 3, and were then variable until day 7 when the virus was cleared. These replication kinetics differ from those we observed for the 1968 H3N2 virus. For the 1968 H3N2 virus, at a DR inoculation dose of 10^6^, viral titers were constant between 1-5 dpi and the virus was cleared between 5-7 dpi. At lower inoculation doses (*i.e.* 10^4^-10^1^) of the 1968 H3N2 virus, viral titers in individual DRs were highly variable. It is unclear why the replication kinetics differed between then 1968 H3N2 virus and the A/Victoria/3/1975 (H3N2) strain. This may be due to differences in ferret sampling procedures or may reflect further mammalian adaptation of the A/Victoria/3/1975 (H3N2) strain resulting from extensive circulation in humans.

To our knowledge, we are the first to assess airborne transmission across decreasing DR inoculation doses in ferrets. Several studies have reported airborne transmission of 2009 pandemic H1N1 virus strains and seasonal H3N2 viruses at high inoculation doses of 10^6^ TCID_50_ or pfu (reviewed in(13)). Our findings of 100% and 62.5% transmission efficiency for the 2009 H1N1 and 1968 H3N2 viruses, respectively, at a DR inoculation dose of 10^6^ TCID_50_ are within the range of transmission efficiency reported in prior studies; although seasonal H3N2 viruses are often reported to transmit to between 66-100% of RCs (13). Importantly, direct contact transmission of seasonal H3N2 viruses has been evaluated at reduced DR inoculation doses in ferrets and guinea pigs (20, 21). In direct contact transmission studies in ferrets, DR ferrets were infected with 10^1^ pfu of the A/Victoria/3/1975 (H3N2) virus and co-housed with a contact animal 24 hours post-infection. In these experiments, all the DRs were productively infected, and they transmitted the virus to 100% of contacts (20). Similarly, when guinea pigs were inoculated with 10^3^, 10^2^, and 10^1^ pfu of the A/Panama/2007/1999 (H3N2) virus and co-housed with contacts, 100% of the contacts became infected (21). In our studies, the 1968 H3N2 virus transmitted to 12.5%, 50%, and 25% of respiratory contacts at low inoculation doses of 10^3^, 10^2^, and 10^1^ TCID_50_, respectively. These discrepancies in transmission are most likely due to airborne transmission imposing a more stringent bottleneck on the number of virions transmitted to contact animals relative to direct contact transmission (22).

Collectively, our studies provide a new approach and metric to study airborne transmission in ferrets. We show that while both the 2009 H1N1 and 1968 H3N2 viruses have similar ID_50_’s, these viruses exhibit differing degrees of transmissibility. For the 2009 H1N1 virus, all infected DRs regardless of inoculation dose transmitted the virus to their RCs, and the ID_50_ and TD50 were equivalent. Moreover, across DR inoculation doses, the 2009 H1N1 virus replicated to similar peak titers and similar amounts of virus were shed. In contrast, for the 1968 H3N2 virus as DR inoculation dose decreased, peak titers and total virus shedding were reduced resulting in reduced transmission and an increased TD_50_. Importantly, the use of TD_50_ has the advantage of permitting comparisons of transmissibility over a log scale. This contrasts the conventional approach of defining transmissibility in 25-33% increments with transmission to greater than 66% of contacts considered efficient. For example, using the conventional high dose inoculation approach, the transmission efficiency of the 1968 H3N2 and 2009 H1N1 viruses would have been 62.5% and 100% versus TD_50_ values of <10^0^ and 10^3.49^, respectively. In the future, it will be valuable to apply the TD_50_ metric in several different contexts. This could include comparing the TD_50_ of seasonal H3N2 viruses collected since 1968 to determine if these viruses have acquired enhanced transmissibility during their circulation in humans. Moreover, it will be valuable to determine the TD_50_ of other pandemic viruses such as representative strains of the 1957 H2N2 and 1918 H1N1 viruses. The TD_50_ of pandemic viruses could then be compared to other zoonotic or emerging viruses that exhibit airborne transmission in ferrets. This may reveal distinct differences in the transmissibility of pandemic viruses relative to zoonotic strains, and this knowledge could then be used to improve pandemic risk assessments.

## Materials and Methods

### Cells and Viruses

Recombinant A/California/07/2009 (H1N1pdm09) and A/Hong Kong/1/1968 (H3N2) viruses were generated by reverse genetics. Reverse genetics plasmids for the A/California/07/2009 (H1N1pdm09) virus were generously provided by Dr. Jesse Bloom, Fred Hutch Cancer Research Center, Seattle, WA. Reverse genetics plasmids for the A/Hong Kong/1/1968 (H3N2) virus were generated via cloning individual gene segments from the A/Hong Kong/1/1968 (H3N2) (mother clone) (NR-28620, BEI Resources). This virus has a defined passage history of 3 monkey cell passages and 3 egg passages prior to plaque purification. Each viral gene was cloned into pDP2000 following previously established methodology, and viruses were rescued in a co-culture of MDCK and 293T cells (23). After virus rescue, the virus was passaged twice in MDCK cells to generate a virus stock. MDCK cells (London Line, FR-58) were obtained through the International Reagent Resource Influenza Division, WHO Collaborating Center for Surveillance, Epidemiology and Control of Influenza, Centers for Disease Control and Prevention, Atlanta, GA, USA. MDCK cells were cultured at 37°C in 5% CO_2_ using DMEM (HyClone) media supplemented with 10% FBS (Seradigm), 4.0 mM L-glutamine (Corning), 15 mM HEPES (Corning), and 1% antibiotic and antimycotic solution (Life Technologies). 293T cells (American Type Culture Collection) were cultured under the same conditions using Opti-MEM media (Invitrogen) supplemented with 10% FBS and 1% antibiotic and antimycotic solution. When propagating virus, the media was replaced with viral culture media consisting of Opti-MEM supplemented with 1% antibiotic and antimycotic solution, and 1 ug/mL of tosylsulfonyl phenylalanyl chloromethyl ketone (TPCK)-trypsin (Worthington).

### Virus Titrations

The tissue culture infectious dose 50% (TCID_50_) of propagated virus stocks was determined on MDCK cells seeded in 24-well plates. Serial 10-fold dilutions of each virus stock in virus culture media were overlayed on four wells of a 24-well plate and then incubated at 37°C for 96 hours. At this time, plates were scored for cytopathic effect (CPE), and the TCID_50_/mL was calculated using the method of Reed and Muench (24). All virus stocks were aliquoted and stored at - 80°C until use. Virus stocks were titrated a minimum of 4 times, and the average viral titer was used to prepare log-fold dilutions of virus for inoculation of ferrets.

For nasal wash samples, the same titration approach was used with the following modifications (16). Nasal wash samples were titrated using a combination of 96 well and 24 well plates of MDCK cells. To determine peak titers, 10-fold serial dilutions of nasal wash were added to a 96-well plate containing MDCK cells in virus culture media. Following inoculation of the MDCK cells, plates were incubated at 37°C and evaluated for CPE after 96 hours. To enhance the limit of detection, 100 uL of nasal wash sample was also added to 2 wells of a 24-well plate and incubated at room temperature for an hour. The media was then replaced with virus culture media, incubated at 37°C for 4 days, and scored for CPE. The TCID_50_/mL of nasal wash was then calculated using the method of Reed and Muench (24).

### Biocontainment and Animal Care and Use

Experiments using recombinant A/California/07/2009 (H1N1pdm09) and A/Hong Kong/1/1968 (H3N2) were performed at biosafety level 2+. All animal studies and procedures were conducted in compliance with all applicable regulations and guidelines. Animal studies were approved by the PSU Animal Care and Use Committee under protocol no. 201800250. Equal numbers of male and female ferrets, age 24-30 weeks of age (Triple F Farms, Sayre, PA) were used for all studies. Two male and two female DR:RC pairs (total 4 DR:RC pairs) were used in each transmission study. All animals were pre-screened by hemagglutination inhibition assay and confirmed seronegative for currently circulating influenza A viruses.

### Ferret Transmission Experiments

To evaluate airborne transmission, donor ferrets (n=4/inoculation dose) were sedated by intramuscular (*i.m.*) injection with a combination of ketamine (30 mg/kg), xylazine (2 mg/kg), and atropine (0.05 mg/kg). Animals were intranasally inoculated with 10 to 100-fold decreasing virus doses (10^6^ – 10^0^ TCID_50_) in a 1 mL volume of virus diluted in Opti-MEM media (Life Technologies, CA). Following inoculation, ferrets were given an intramuscular (*i.m.*) injection of the reversal agent, atipamezole (0.5 mg/kg), and were housed in individual biocontainment ferret cages (Allentown, PA). Twenty-four hours post-inoculation, each inoculated DR ferret was placed on one side of a transmission cage and paired with a RC ferret. Transmission experiments were performed using large stainless steel ventilated ferret cages (Allentown, PA) modified such that DR and RC pairs were separated by a 5 cm wide perforated offset divider.

At the time of introduction into transmission cages, and every other day for 14 days, DR ferrets were sedated by *i.m.* injection with ketamine (20 mg/kg), xylazine (2 mg/kg), and atropine (0.05 mg/kg), and nasal wash samples were collected by instilling a 1 mL volume of PBS into the nose and inducing sneezing onto a petri dish. An additional 1 mL of PBS was then used to rinse the dish and nasal wash samples were aliquoted and stored at -80°C. The RC ferrets were sampled on alternating days from the DRs, and nasal wash samples were collected following the same procedure. All animals were monitored daily for clinical signs of illness, and body weight and temperatures were recorded at the time of nasal wash. At 21 dpi of DRs, all ferrets were deeply sedated. Blood was then collected via cardiac puncture, and animals were euthanasia via overdose with sodium pentobarbital. Blood was processed to recover serum, which was stored at -20°C.

### Serology

Serum samples were evaluated for antibody titers using a hemagglutination inhibition (HI) assay. Serum was treated with receptor-destroying enzyme (Hardy Diagnostics) overnight, followed by heat inactivation at 56°C for 1 hour and further diluted to 1:10 with normal saline. Sera was then diluted 2-fold in PBS in 96-well V-bottom plates (Corning), incubated with 8 HA units of virus/well, and then overlayed with a 0.5% suspension of turkey red blood cells (Lampire Biological Laboratories). After 35-40 minutes, hemagglutination was visually assessed, and the HI titer was determined as the reciprocal of the lowest serum dilution without agglutination.

### Statistical Analyses

ID_50_ and TD_50_ were calculated using the method of Reed and Muench (24). Area under the curve analyses to assess total viral shedding, and comparisons of peak nasal wash titers were performed using GraphPad Prism (version 10.2.1). For comparisons between DR inoculation doses for a specific virus strain, Kruskal-Wallis tests were performed with the two-stage step-up procedure of Benjamini, Krieger, and Yekutieli, to control for False Discovery Rate. For comparisons between virus strains at a specific dose, Mann-Whitney U tests were performed. For all statistical analysis p<0.05 was considered significant.

## Acknowledgments

CJF, KMS, DRP, and TCS planned and performed all experiments with assistance from VCW and DGS. CJF and TCS drafted and edited the manuscript. All authors have no competing interests. We acknowledge the Pennsylvania State University Animal Resource Program staff for their support with animal studies. This project has been funded in whole or in part with Federal funds from the National Institute of Allergy and Infectious Diseases, National Institutes of Health, Department of Health and Human Services, under Contract No. 75N93021C00017 (NIAID Centers of Excellence for Influenza Research and Response, CEIRR). Support was also provided by the USDA National Institute of Food and Agriculture, Hatch project 4955 (TCS).

